# Westwards expansion of the European catfish *Silurus glanis* in the Douro River (Portugal)

**DOI:** 10.1101/2023.01.07.522915

**Authors:** Christos Gkenas, Joana Martelo, Diogo Ribeiro, João Gago, Gil Santos, Diogo Dias, Filipe Ribeiro

## Abstract

The current study reports the first occurrence and the spread of the European catfish *Silurus glanis* (Family: Siluridae) in the Portuguese section of the Douro River, suggesting a potential expansion of its distribution in Portugal either via westward dispersal across international rivers and/or human-assisted introductions into new reservoirs and drainages. European catfish has unique features (e.g., opportunistic predator, hunting, and aggregation behaviour) that make it highly suitable for establishing self-sustaining populations in new areas and likely contribute to its invasion success. The species may severely affect native prey communities and modify food web structure and ecosystem functioning. Efficient and sustainable management actions are needed to prevent further introductions in the future.

## INTRODUCTION

Non-native species have been identified as a leading threat to aquatic biodiversity worldwide (Sala et al., 2000), being implicated in biota declines, ecosystem degradation and ecosystem service changes (Rahel & Olden, 2008; Reid et al., 2018). The most common vectors for the introduction of non-native fish (NNF) in Europe are aquaculture, recreational fisheries and ornamental fish trade (Gozlan et al., 2010), though their relevance varies geographically. For instance, while aquaculture is a main driver of NNF introduction and establishment in central Europe, recreational fisheries predominate in the Iberian Peninsula (Ribeiro et al., 2009; Carpio et al., 2019). This region is a freshwater fish invasion hotspot (Leprieur et al., 2008), with one new species being recorded every two years (Ribeiro et al., 2009; Banha et al., 2017). Most introductions of NNF are associated with recreational angling, with introduced fish being either used as live baits or as fishing targets (Banha & Anastácio, 2015; Carpio et al., 2019), and some large apex predators being highly valued as trophies (Gago et al., 2016; Ribeiro et al., 2021). Introductions of apex predators may have major impacts on native communities and food web structure (Cucherousset et al., 2018), which can be magnified in Iberian rivers with high endemism of small-bodied endangered fish and absence of native piscivorous fishes (Copp et al., 2009).

European catfish, *Silurus glanis*, represents one of the most successful and concerning fish introductions in southern and western Europe (Copp et al., 2009; Cucherousset et al., 2018). Native to eastern Europe and western Asia, the species began to disperse across European river basins in the first half of the 19^th^ century, and it is now very widespread in at least 13 European countries (Castaldelli et al., 2013; Boulêtreau & Santoul, 2016; Vejřík et al., 2017; Rees et al., 2017), including Spain and Portugal (Benejam et al., 2007; Gkenas et al., 2015). Mainly introduced for trophy angling, European catfish is an extremely large top predator displaying opportunistic feeding behaviour, and as such is expected to have highly detrimental effects on recipient native communities (Copp et al., 2009; Cucherousset et al., 2018). Although there is limited knowledge on European catfish impact in the Iberia Peninsula (e.g. Ferreira et al., 2019), the species has been reported elsewhere to threaten not only native fish populations (Guillerault et al., 2017), but also birds, amphibians and mammals (Cucherousset et al., 2012; Boulêtreau et al., 2018; 2021), and to potentially compete with native top predators, such as pike *Esox lucius* (Vejřík et al., 2017). Management and control actions for European catfish are therefore needed, and an important first step forward is to better understand and characterise the species range expansion, particularly at an early invasion stage (Green & Grosholz, 2021).

Here, we report the first scientific record of the invasive European catfish in the Douro River (Portugal) and provide an update on citizen science data documenting the spread of catfish in the Portuguese Douro River. We further present results of the diet of a single catfish and discuss the potential impacts of the species establishment on riverine fish populations in the Douro River basin.

## MATERIALS AND METHODS

A single specimen of *S. glanis* was collected by a fisherman using gillnets, in Carrapatelo reservoir (41°06’53.5”N 7°57’56.7”W), mainstem of Douro River, on March 12, 2022 and an additional citizen science record was reported by a fisherman in Régua reservoir in 2022 (Fig. 1; Table S1 (see Supplementary information, available at http://www.limnetica.net/en/limnetica)). After collection, the specimen was donated and transported to the laboratory for identification following Kottelat and Freyhof (2007). Measurements of total length (TL ± 1 mm), standard length (SL ± 1 mm) and total weight (TW ± 0.1 g) were determined. The catfish was dissected to examine the stomach for diet analysis and to determine sex through macroscopic evaluation of the gonads. Stomach content was identified to the lowest possible taxonomic level and prey were recorded. The specimen was photographed in their natural condition (Fig. 2) and deposited in the zoological collection “Museu Bocage” of the Museu Nacional de História Natural e da Ciência (MUHNAC; Lisbon, Portugal) with the accession number MB05-3635.

**Figure 1.**
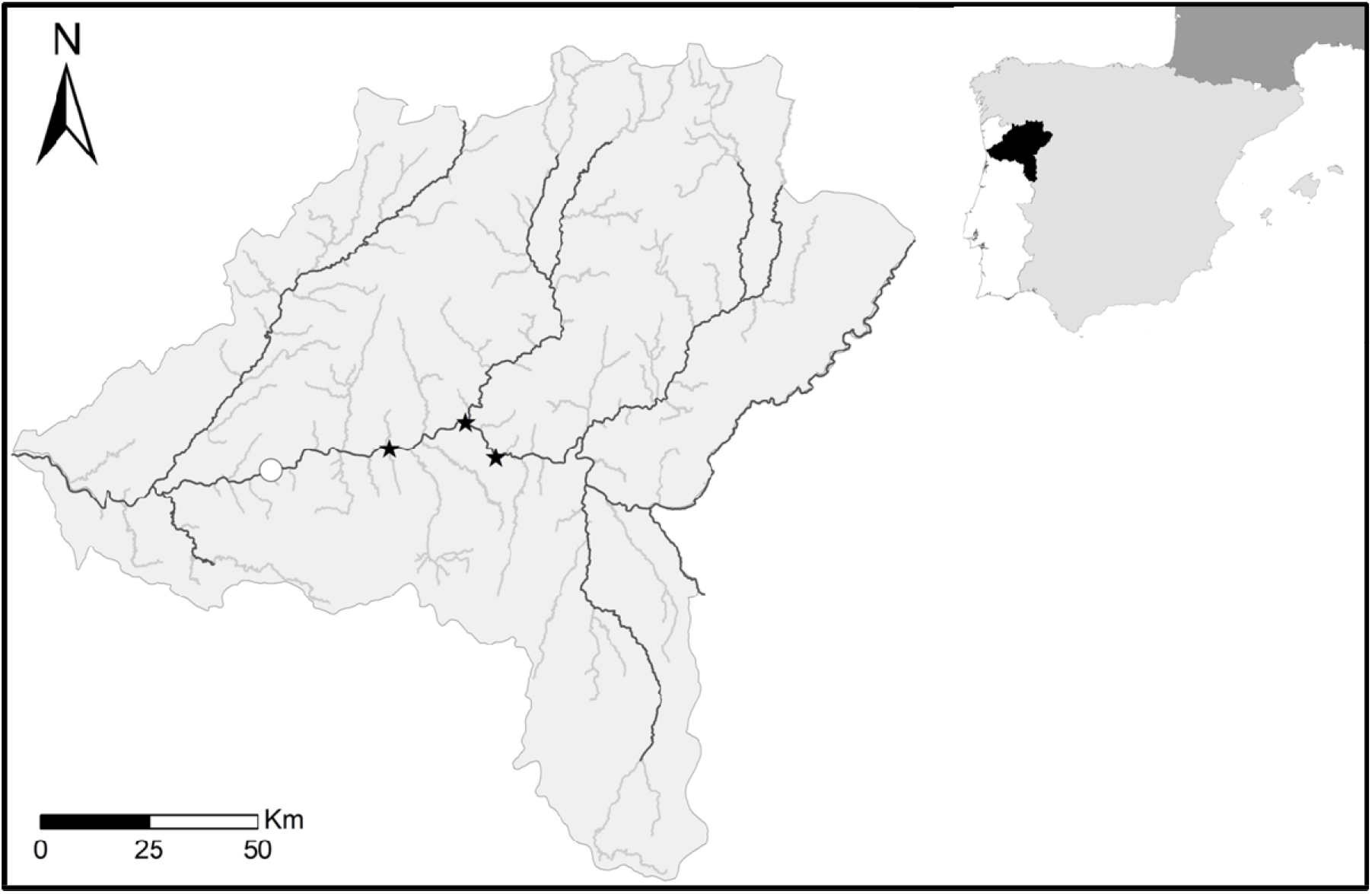
Map of *Silurus glanis* occurrences in the Portuguese part of the Douro River. Records depict scientific (white circle) and citizen science data (black star).

**Figure 2.**
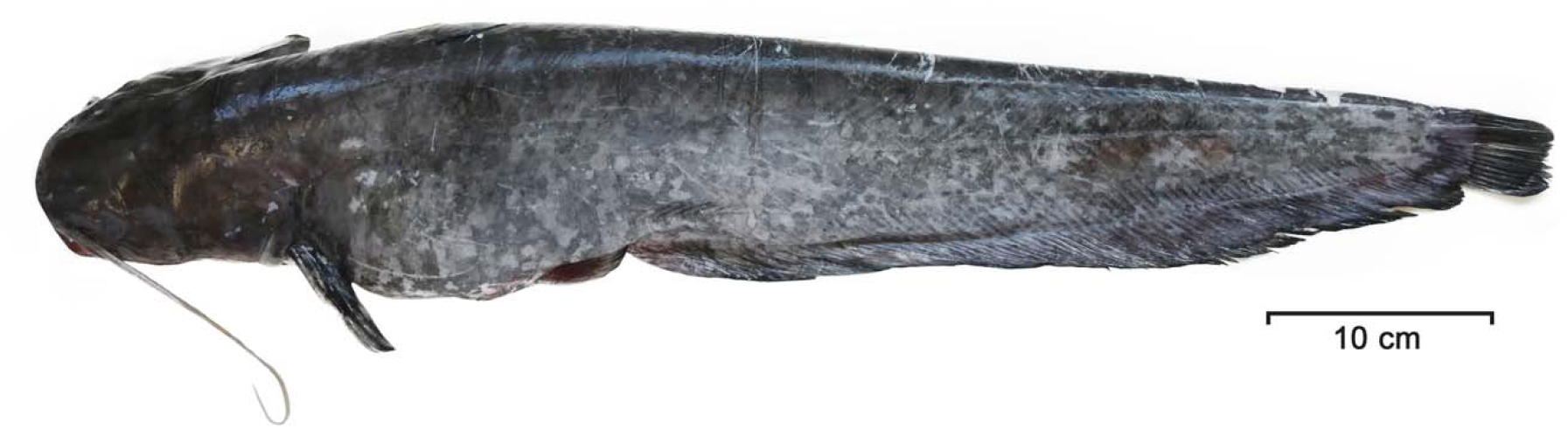
Specimen of *Silurus glanis* (690□mm total length and 1920□g) captured in the Douro river (Carrapatelo reservoir, Portugal) and deposited in the zoological collections “Museu Bocage” of the Museu Nacional de História Natural e da Ciência (MUHNAC; Lisbon, Portugal).

## RESULTS

The collected specimen was a male, measured 690 mm (TL), 620 mm (SL) and weighed 1.920 kg (TW). Based on the age data provided by Copp et al. (2009) and Alp et al. (2011), the specimen had an approximate age of 2-3 years. The body was elongated, laterally compressed after the head, and the skin was smooth and scaleless. The head was broad and flat with small eyes, widely spaced nostrils and a large mouth containing lines of numerous small teeth. Two long and slender barbels on the upper jaw and four short flexible barbels below the lower jaw were present. The colour of the body was dark along the back with lighted sides and a white abdomen. Morphometric and meristic data corroborated the identification, displaying 5 dorsal finrays, I, 12 ventral finrays, I, 12 pectoral finrays, 83 anal finrays and 18 caudal finrays. Stomach content analysis revealed two freshwater shrimps (*Atyaephyra desmarestii* L.) and the remains of an unidentified aquatic insect.

## DISCUSSION

In recent decades, *S. glanis* has been increasingly introduced for recreational angling owing to demand for large-bodied specimens by trophy anglers, as their addition to freshwater fisheries is likely to boost annual revenue (Rees et al., 2017). Since its first record to the Iberian Peninsula in 1974 at Mequinenza-Ribarroja reservoir (Ebro drainage, Doadrio, 2002), European catfish has been recorded across Catalonia (Benejam et al., 2007), Tagus (Pérez-Bote & Roso, 2009) and Guadalquivir drainages (Moreno-Valcárcel et al., 2013; Fernández-Delgado et al., 2014; Sáez-Gómez and Prenda 2019). Catfish has recently reached Portugal, probably through downstream dispersal along the Tagus River from Spain (Gkenas et al., 2015), which suggests that this species follows a westward expansion in the Iberian Peninsula. However, fishermen translocations cannot be ruled out, given that this has been the main vector of fish introductions into new drainages (Anastácio et al., 2018). Our study updates this distribution, by documenting the presence of European catfish in the Carrapatelo reservoir (Douro mainstem, Portugal) which is about 60 km far from the river mouth, thus it is likely that the species also follows a west dispersal, either natural or human-assisted dispersion, and possible spread to the lower Douro River. Its first record in the Douro River was 8 years earlier (2014) in the surroundings of Soria (Spain, Eastern Iberian Peninsula) (Parrondo et al., 2018). Since then it has spread rapidly in Portugal, being subsequently detected by anglers in Valeira reservoir in 2019 and in Régua reservoir in 2020 and 2022 (Fig. 1; Table S1; Martelo et al., 2021), suggesting an approximate annual dispersion of about 30 km/yr. Considering the potential age of our single specimen relative to the size, the arrival to the Carrapatelo reservoir of European catfish happened approximately 2 to 3 years ago (c.a. 2019-2020), suggesting that this apex predator could have already invaded the entire mainstem of the Douro River.

Nevertheless, this study was constrained by low number of records, with a single capture not providing sufficient information to determine if there is an already established population in the Douro River. However, its establishment is highly likely given that previous (Parrondo et al., 2018; Martelo et al., 2021) and current catfish records observed in the Douro River represent a similar spatial pattern and range expansion of the European catfish found in the Tagus River drainage, where it is thriving (Gkenas et al., 2015; Gago et al., 2016). Additionally, the inclusion of a few citizen science records may not provide a representative sample, thus making it difficult to draw reliable conclusions about its current distribution, though fishermen records are increasingly being used successfully to describe the range expansion of invasive species (Poursanidis & Zenetos, 2013; Morais et al., 2017). Therefore, it is important to use caution when interpreting sporadic records and consider collecting scientific data over a longer period of time in order to verify the species expansion in the Douro River.

The ability of European catfish to colonise novel ecosystems seems to be driven by its wide diet plasticity and adaptability to newly available prey (Syväranta et al., 2010; Cucherousset et al., 2012). Our preliminary stomach content analysis comprised only three items (2 freshwater shrimps and 1 unidentified insect), suggesting predation close to river banks where *A. desmarestii* are numerous (Fidalgo & Gerhardt, 2002) and the use of shallow habitats during catfish’s foraging activity (Santos 2022). This needs further analysis given our low sample size, but is consistent with previous work (Ferreira et al., 2019). In addition, we observed low stomach fullness, which might be a result of decreased food availability (Vejřík et al., 2017), long permanence of catfish in the nets (de Santis & Volta, 2021), or regurgitation on capture (Guillerault et al., 2017). In line with previous studies of catfish diet (e.g., Ferreira et al., 2019; de Santis & Volta, 2021), we expect an increase in piscivory as the catfish population in the Douro River becomes older, given that larger individuals are expected to feed on larger preys to meet their dietary needs (de Santis & Volta, 2021). Furthermore, the arrival of European catfish to the Douro River may impact populations of migratory Atlantic salmon, *Salmo salar*, particularly those in the lower section of the river, which is the southern distribution limit of the species (Godinho & Pinheiro, 2019). This raises concerns for the recent efforts to improve river’s connectivity, once European catfish often targets migratory fish while the latter are negotiating fish passages or during spawning aggregations (Boulêtreau et al., 2018, 2021).

Catfish display unique behaviours in the invaded range, such as massive aggregations in groups (Boulêtreau et al., 2011) and beaching behaviour (Cucherousset et al., 2012), that may potentially have various impacts on Douro River native communities, including nutrient translocation from catfish feeding areas, hence subsequently affecting primary production, nutrient cycling and predator-prey interactions (Boulêtreau et al., 2011). Additionally, predation on small mammals and waterfowls often exhibited by catfish may have unexpected implications on consumer-resources dynamics and ultimately on ecosystem functioning (Cucherousset et al., 2018; Milardi et al., 2022).

Highlighting that the presence of *S. glanis* in the mainstem of Douro River was probable due to natural dispersal but also to deliberate releases or unregulated transfer activities by sport anglers, there is the necessity of doing educational awareness campaigns to the general population and fisheries associations about the threats to biodiversity posed by catfish’s dispersal into the wild (Rees et al., 2017). In addition, scientific divulgation could disseminate knowledge to society about the consequences of introducing a non□native species in a new environment, and the importance of preserving the native communities. Therefore, more comprehensive studies concerning the species distribution and impact are crucial to guide management and control actions which should involve decision-makers and key stakeholders, such as fishermen, to prevent further invasion and contribute to early warning records.

## ACKNOWLEDGEMENTS

We are thankful to the recreational fishermen who voluntarily donated the specimen and provided georeferenced data on the European catfish and Miguel Santos for improvement on the image of the catfish.

## SUPPLEMENTARY MATERIAL

Table S1. Geo-referenced records of *Silurus glanis* in the Douro River.

